# Induced pluripotent stem cells and cerebral organoids from the critically endangered Sumatran rhinoceros

**DOI:** 10.1101/2022.05.12.491654

**Authors:** Vera Zywitza, Silke Frahm, Norman Krüger, Anja Weise, Frank Göritz, Robert Hermes, Susanne Holtze, Silvia Colleoni, Cesare Galli, Micha Drukker, Thomas B. Hildebrandt, Sebastian Diecke

**Author notes:** Correspondence (S.D.).

## Abstract

Less than 80 Sumatran rhinos (SR, *Dicerorhinus sumatrensis)* are left on earth. Habitat loss and limited breeding possibilities are the greatest threats for the species and lead to a continuous population decline. To stop erosion of genetic diversity, reintroduction of genetic material is indispensable. However, as the propagation rate of captive breeding is far too low, innovative technologies have to be developed. Induced pluripotent stem cells (iPSCs) are a powerful tool to fight extinction. They give rise to each cell within the body including gametes, and provide a unique modality to preserve genetic material across time. Additionally, they enable studying species-specific developmental processes.

Here, we generate iPSCs from the last male Malaysian SR Kertam, who died in 2019, and characterize them comprehensively. Differentiation in cells of the three germ layers and cerebral organoids demonstrate their high quality and great potential for supporting rescue of this critically endangered species.

**HIGHLIGHTS:**

- Characterization of Sumatran Rhino (SR) fibroblasts
- Generation of SR induced pluripotent stem cells (SR-iPSCs)
- SR-iPSCs generate cells of the three germ layers
- SR-iPSCs give rise to cerebral organoids

## INTRODUCTION

The Sumatran rhino (SR, *Dicerorhinus sumatrensis)*, also known as hairy or Asian two-horned rhino, is the smallest and most ancient of five extant rhinoceros species (Liu et al., 2021). Once, it inhabited a continuous, vast area in East and Southeast Asia, but now only small, fragmented populations remain scattered across Sumatra and Indonesian Borneo. The current stock is estimated at 40-78 individuals distributed over up to ten subpopulations (Emslie et al., 2019), which increases the risks of inbreeding and asymmetric reproductive ageing related female infertility, i.e. development of reproductive tract pathologies as a result of repeated ovarian cycling activity due to absence of naturally occurring long acyclic gestation and lactation phases (Hermes et al., 2004). Reintroduction of genetic material is crucial to support a healthy and self-sustainable SR population. Contrary, the classical captive breeding approach in the USA, Malaysia and Indonesia only led to five offspring from a total of 47 individuals since the first SRs were received in captivity for breeding purposes in 1985 (Sumatran Rhino Crisis Summit, Singapore Zoo, 1-4 April 2013) highlighting the need for future application of advanced assisted reproduction technologies (Hildebrandt et al., 2021). Toward this aim, we will employ adult, somatic cells to generate induced pluripotent stem cells (iPSCs) (Takahashi et al., 2007), which - in contrast to primary cells - are capable of unlimited expansion and therefore a valuable tool to store genetic information of endangered species across time. Additionally, iPSCs can make each cell type of a body including oocytes and spermatozoa (Hayashi et al., 2021), which in turn could be used for *in vitro* fertilization thereby contributing to breeding of infertile or already deceased individuals (Hildebrandt et al., 2021; Saragusty et al., 2016). Beyond application in innovative conservation strategies, iPSCs from endangered species enable studying species-specific developmental processes. As embryonic material from exotic, large mammals is very limited to almost inaccessible, iPSCs are an unprecedented tool to gain insights into embryo- and organogenesis.

Here, we generate iPSCs from the SR Kertam, who was the last male Malaysian SR before his death at Tabin Wildlife Reserve in 2019. We characterize both source fibroblasts and iPSCs thoroughly, and demonstrate three-germ layer differentiation potential of the latter.

Generation of cerebral organoids from SR-iPSCs highlights their ability to generate complex 3D structures, and concomitantly represents a promising application for studying evolutionary progression of brain development across species. Taken together, this work represents the first step towards fighting extinction of the SR using stem cell associated techniques (SCAT).

## RESULTS

### Generation and Characterization of SR-iPSCs

In 2019, the Sumatran rhino (SR) was declared extinct in Malaysia. The last three individuals, Puntung, Imam (both female), and Kertam (male) were housed at Tabin Wildlife Reserve in Lahad Datu, Malaysia, and great efforts have been made to breed them in captivity. Even after Kertam’s death, an *in vitro* fertilization attempt using his cryopreserved sperm has been made. However, Imam’s isolated oocytes failed to divide after fertilization, probably due to poor quality of Kertam’s cryopreserved spermatozoa. To preserve the genetic material and to retain the possibility to obtain viable sperm cells from Kertam in future, we generated induced pluripotent stem cells (iPSCs). Therefore, we took a skin biopsy under general anesthesia and isolated fibroblasts. Cryopreserved SR-fibroblasts grew well after thawing, and had a population doubling time of 19 hours during the logarithmic phase of growth (48-96 hours, Figure 1B). To confirm their identity, we sequenced a 230 bp cytochrome-b region, which can be used to discriminate the five extant rhinoceros species (Ewart et al., 2018). Except for one substitution (C → T), the amplified DNA mapped unambiguously to the SR-reference sequence (Figure 1A). To generate iPSCs, we used an adapted protocol that has been optimized recently for reprogramming northern white rhino (NWR, *Ceratotherium simum cottoni*) fibroblasts (Hayashi et al., *submitted*; Korody et al., 2021; Figure 1C). Therefore, SR-fibroblasts were plated at low density in skin media and two days later transduced with Sendai-Viruses encoding the human reprogramming factors *POU5F1, SOX2, KLF4* and *c-MYC*. Similar to NWR-fibroblasts, SR-fibroblasts were less susceptible to viral transduction than human fibroblasts. A two times higher viral concentration as recommended for human cells worked well (Figure 1 D, day 7).

**Figure 1.**
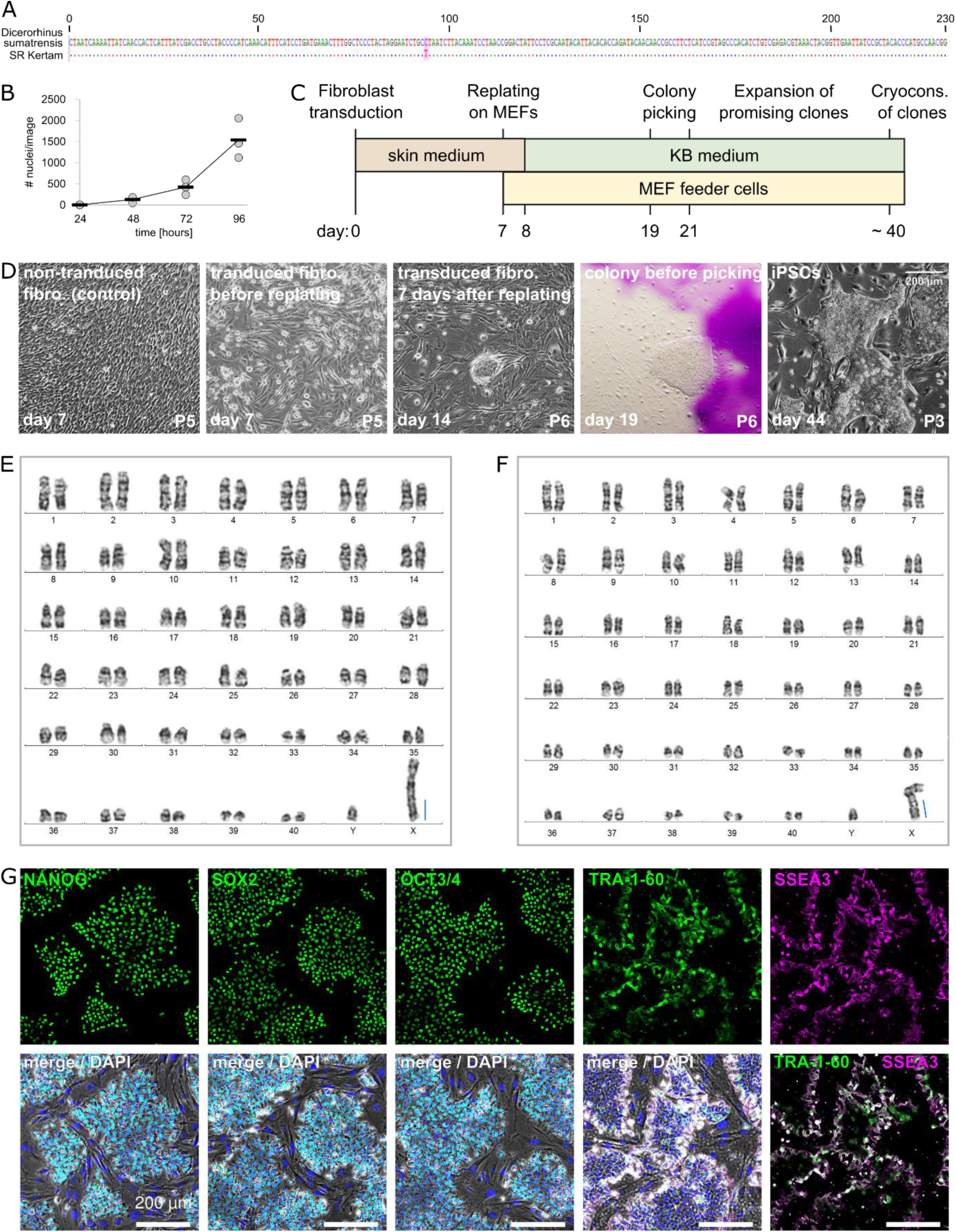
Generation of SR-iPSCs. See also Supplementary Figure 1. **A**. Species analysis based on amplification and Sanger-sequencing of a 230 bp cytochrome-b region, according to Ewart et al., 2018. Top line: published sequence. Substitution is highlighted. **B**. Proliferation analysis. Stained nuclei of cells fixed at 24h, 48h, 72h and 96h after plating were counted in three images per timepoint (one per 24 well). The slope between 48h and 96h was used to determine the population doubling time. **C**. Timeline of reprogramming. **D**. Cell images during reprogramming of SR Kertam fibroblasts (10x magnification). Scale bar image far right: 200 μm. **E. and F**. Exemplarily G-banded karyotypes of SR Kertam fibroblasts (E) and iPSCs (F). 2n = 82, including Y and X chromosome. Blue bar indicates addition of heterochromatin near the q terminal ends of the X chromosome. **G**. Immunostaining of pluripotency markers. Upper lane: marker only. Lower lane: merge marker, phase contrast and DAPI. Bottom far right: merge TRA-1-60 and SSEA3. Scale bars: 200 μm.

Within 7 days after transduction, the characteristic mesenchymal to epithelial transition (MET) became apparent, which was accompanied by morphological transformation from spindle- to planar-shaped with high nucleus to cytoplasm ratio. On day 7, cells were replated on mitotically inactivated mouse embryonic fibroblasts (MEFs) and feeder-free on Geltrex-coated dishes. The day after, cultures on MEFs were changed to KB medium, whereas feeder-free cultures were changed to mTeSR1. Individual colonies of tightly-packed cells with prominent nucleoli were observed from 14 days after transduction on, and were isolated from both conditions on days 19 and 21. Similar to reprogramming of NWR-iPSCs, SR-iPSC lines could only be established from KB on MEFs conditions. However, once stabilized, NWR-iPSC lines could be adapted to feeder-free mTeSR1 (Korody et al., 2021) or rather mTeSR1:KB medium (Hayashi et al., *submitted*), whereas SR-iPSCs spontaneously differentiated or died in all tested feeder-free conditions. To assess the genetic integrity of SR-fibroblasts and SR-iPSCs, we performed karyotyping by GTG banding and observed a diploid chromosome number of 2n = 82 comprising 80 acrocentric autosomes and two gonosomes (X and Y) in both cell types. As described previously (Houck et al., 1995), the X chromosome contained a conspicuous long q arm (blue bar Figures 1 E and F), which demonstrably consists of heterochromatin (CGB banding, Supplementary Figure 1). SR-iPSCs expressed the canonical pluripotency markers NANOG, SOX2, OCT3/4 as well as the pluripotency surface markers TRA-1-60 and SSEA3 (Figure 1G).

### Three germ-layer differentiation potential of SR-iPSCs

Next, we demonstrated the pluripotent potential of SR-iPSCs by generating cells of the three germ layers using protocols optimized for directed differentiation of human PSCs. Definitive endoderm progenitors were efficiently generated after 5 days of culture in differentiation medium, and their identity confirmed by immunostaining of GATA4, GATA6 and SOX17 (Figure2A). To represent the mesoderm, SR-iPSCs were differentiated into beating cardiomyocytes. Clusters of beating cells were observed 9 days after start of differentiation.

The cells expressed the cardiac specific proteins ACTN2 and TNNT2 (Figure 2B). Towards ectoderm, we differentiated SR-iPSCs into neural progenitor cells and at day 12 of differentiation observed the formation of neural rosettes expressing SOX2, NES, and SOX1 (Figure 2C).

**Figure 2.**
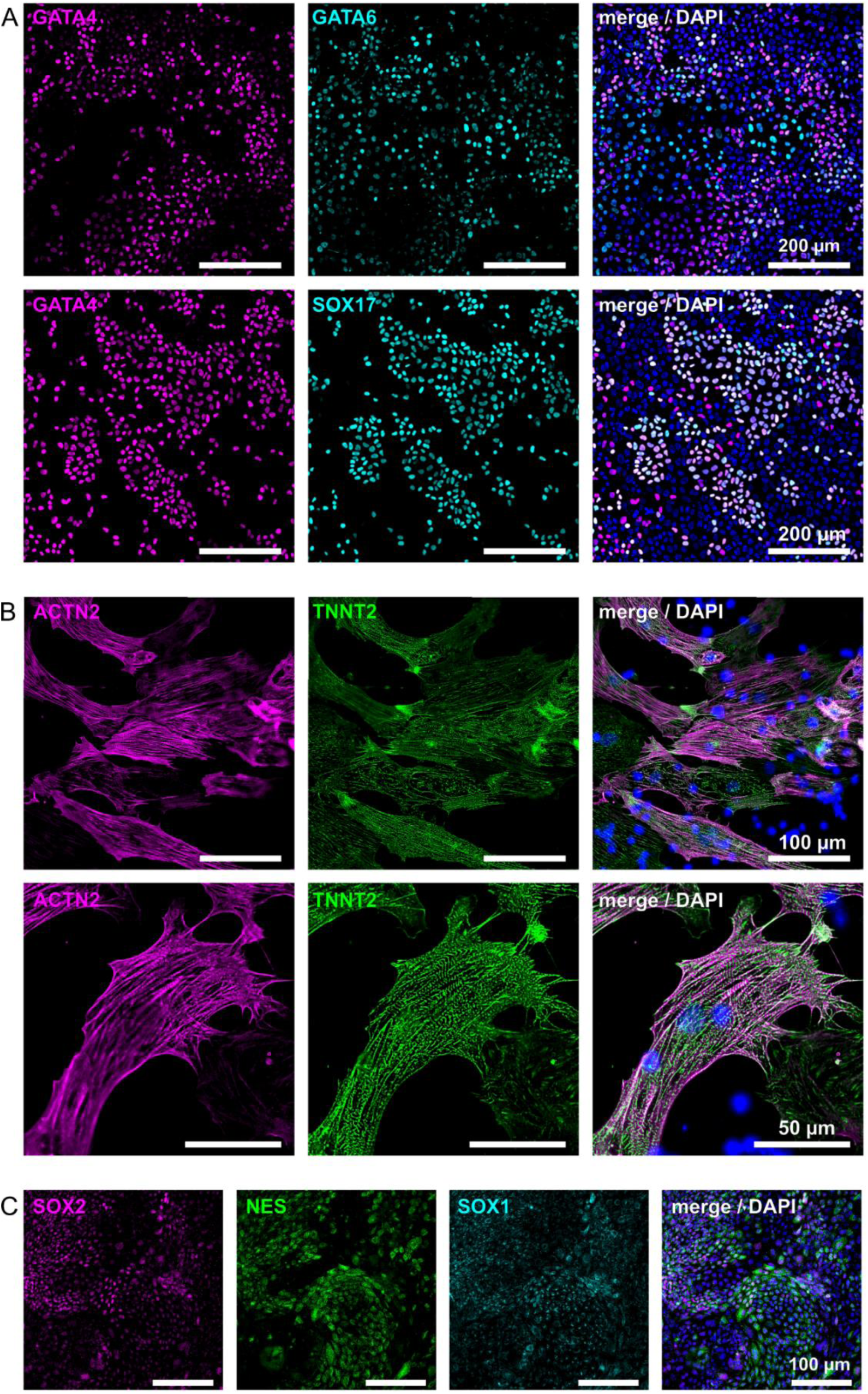
SR-iPSCs generate cells of the three germ layers. Representative immunostainings for markers of the three germ layers upon differentiation of SR-iPSCs towards endoderm (**A**), cardiomyocytes (mesoderm, **B**), and neural precursors (ectoderm, **C**).

### SR cerebral organoids

Little is known about the rhino brain and its development during embryogenesis. In 1878, the morphology of the SR brain has been described (Garrod, 1878) and it seems that not much more is presently known about the brain of the species. The main reason for this lack of knowledge is probably the absence of biological material that can be studied. Much of our understanding of human-specific neural development arose in the last years, since the generation of three-dimensional cerebral organoids from great ape iPSCs opened entirely new possibilities to study early brain expansion and neurogenesis (Benito-Kwiecinski et al., 2021; Lancaster and Knoblich, 2014). Besides great apes, only mouse brain organoids have been generated from embryonic stem cells so far (Eiraku et al., 2008; Nasu et al., 2012). These recapitulated the spatial and temporal aspects of *in vivo* rodent corticogenesis, validating organoids as tools to study brain development. Thus, generation of SR cerebral organoids opens unprecedented possibilities to study rhino neurogenesis.

Having established the neural induction of SR-iPSCs (Figure 2C) using small molecules designed for human NPC differentiation, we thought to employ a standard protocol for cerebral organoid generation (Lancaster and Knoblich, 2014). Standard single-cell seeding did not result in the formation of embryoid bodies (EBs) containing neuroectodermal structures from SR-iPSCs (not shown) in contrast to human iPS cells (Supplementary Figure 2A). Therefore, we modified the protocol and exchanged the enzymatic single-cell seeding at day 0 for a cluster split using a non-enzymatic dissociation procedure. The resulting small aggregates of variable size (100-300 μm in diameter) resembled EBs and developed optically translucent tissue at the margin between days 5 and 7, indicating induction of neuroectoderm (Figure 3A). Further addition of 2% Matrigel to the medium gave rise to radially organized neuroepithelium (a hallmark of human brain organoids) in both, rhino and human organoids (days 10-13; Figure 3 and Supplementary Figure 2). Immunostaining of SR brain organoids at 1 month of age confirmed the formation of well-defined progenitor zones expressing SOX2, some of which were positive for the dorsal cortical marker PAX6 (Figure 3B). Accordingly, SR organoid tissue displayed neurons distal to the progenitor zones which stained positive for the pan-neuronal marker MAP2, the cortical neuronal markers BRN2 and CTIP2, and the hippocampal marker PROX1 (Figure 3B), comparably to human organoids generated with the same kit (Supplementary Figure 2B). After additional 2 months in culture, we observed GFAP-positive astrocytes lining the outer surface of the organoids, as described before in self-organized human cerebral organoids (Renner et al., 2017). Moreover, 3-month old in comparison to 1-month old organoids exhibited more CTIP2-positive neurons, which were lined by SATB2-positive neurons, indicating further maturation.

**Figure 3.**
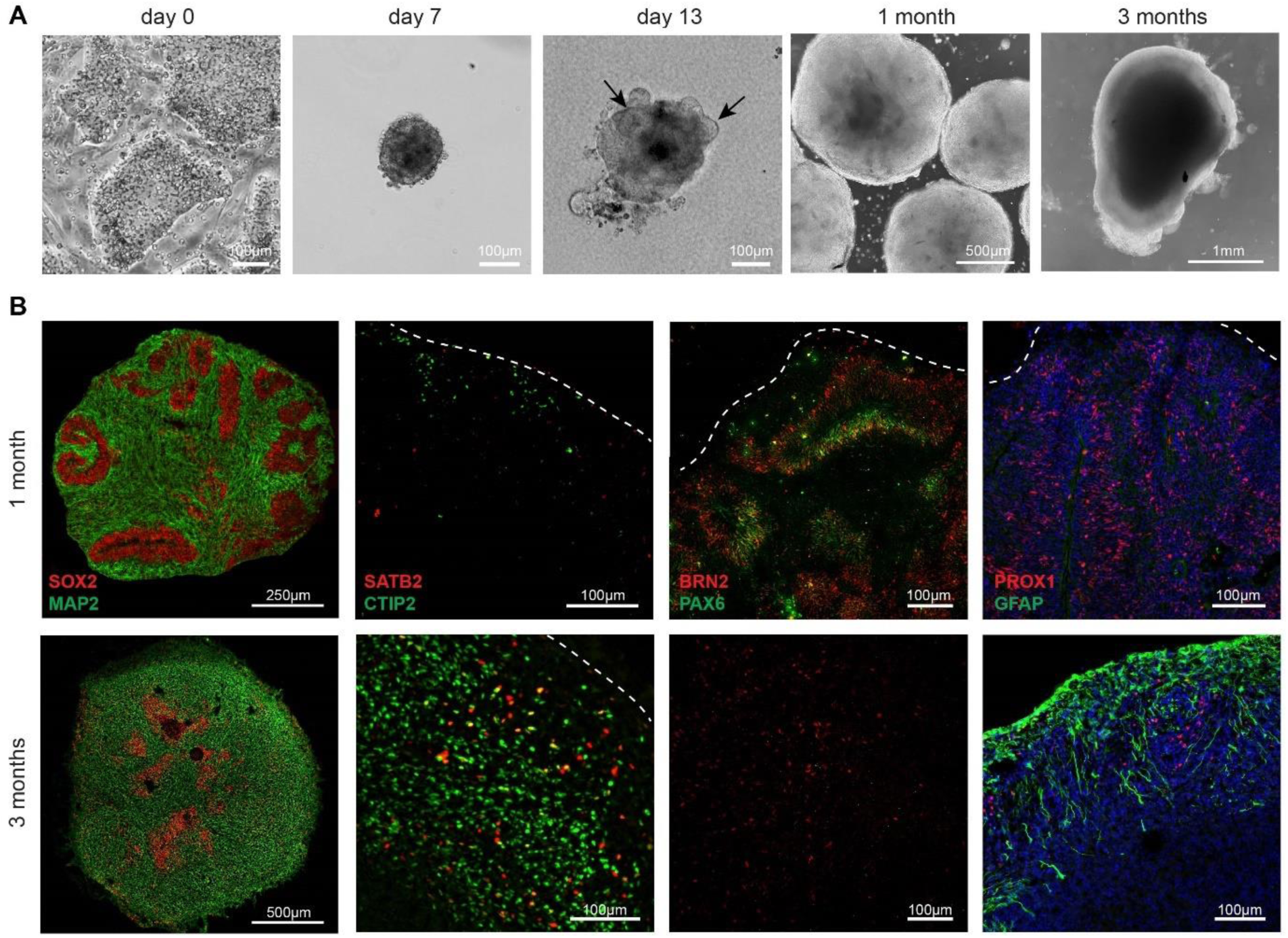
SR cerebral organoids. See also Supplementary Figure 2. **A**. Exemplary phase contrast images during organoid generation at corresponding days of differentiation. Day 0: feeder-dependent SR-iPSC colonies before dissociation; day 7: embryoid body showing a smooth and brightened surface; day 13: after addition of Matrigel to the medium, organoids showed neuroepithelial bud outgrowth resembling neural tube like structures (arrows), which are still visible after 1 month in culture. 3 months: mature organoid. Scale bars are defined within each image. **B**. Immunostaining of 1 (top panels) and 3 (lower panels) months old SR brain organoids. Radially organized neural progenitors stain positive for SOX2 and PAX6, while neurons express MAP2, CTIP2, SATB2, BRN2 and PROX1. Astrocytes appear in older organoids and express GFAP. Scale bars are defined within each image.

## DISCUSSION

The sixth mass extinction, which is largely caused by human activities, is progressing with unprecedented speed (Ceballos et al., 2020; Kolbert, 2015). Notably, the five extant rhinoceros species are particularly affected due to poaching, habitat destruction and fragmentation. Loss of keystone species, especially megavertebrates, can initiate so called “vortex effects” (Lacy, 1993), i. e. accelerated loss of species or entire species societies whose life histories are directly or indirectly connected to the keystone species. The Sumatran Rhino (SR) is known to play a key role in shaping forests and spreading seeds of at least 79 different plant species (McConkey et al., 2022). Many of those evolved alongside megafauna and are designed to be eaten and dispersed by large animals such as elephants and rhinos. Given the differences in ranging and foraging behavior of SRs and elephants, and the fact that 35% of the identified megafaunal-fruits are only distributed by SRs (McConkey et al., 2022), the composition and ecological balance of the Asian rainforest seems highly depending on the SR and loss of the species could have catastrophic consequences.

To fight extinction of the SR, great efforts worldwide - particularly by the Indonesian government and international partners - are made to increase its population. Recently, on March 24, 2022, the rare event of a SR calf born in captivity has been reported by Indonesia’s Ministry of Environment. However, the number of both fertile individuals and successful breeding in captivity remains low and highlights the need to explore innovative alternatives. Stem cell associated techniques (SCAT) in combination with advanced assisted reproduction technologies (aART) open up new perspectives to rescue endangered species (Hildebrandt et al., 2021). By generating induced pluripotent stem cells (iPSCs) from the SR Kertam, who was the last male Malaysian SR and belonged to the Bornean subspecies, we conserved his genetic information, and created an opportunity to produce viable spermatozoa for breeding purposes in future. As the quality of semen collected from SRs is poor directly after retrieval and even worse after cryopreservation and thawing (Agil et al., 2004), *in vitro* generated spermatozoa offer a great alternative for assisted breeding of SRs in general.

In addition, we demonstrate the applicability of iPSCs obtained from endangered species to study species-specific developmental processes by differentiating cerebral organoids from SR-iPSCs. Our experiments show that SR cerebral organoids develop in a self-organized manner, and express the same neural markers as human iPSC-derived brain organoids. For example, SATB2 was detected in neurons of matured organoids. Notably, the timing of SATB2 transcriptional activity defines axonal projection routes exclusive in eutherian mammals, which is relevant for the evolution and development of cortical circuits (Paolino et al., 2020). Previous work with brain organoids from different species has shown that differential temporal expression dynamics of other conserved transcription factors can also have a profound impact on brain expansion (Benito-Kwiecinski et al., 2021). Thus, using comparative RNA-sequencing, SR brain organoids could contribute to study evolutionary progression of brain development in mammals beyond mouse/human interspecies differences and may help to unravel the ancient history of the rhinoceros family.

## EXPERIMENTAL PROCEDURES

### RESOURCE AVAILABILITY

#### Lead Contact

Further information and requests for resources and reagents should be directed to and will be fulfilled by the lead contact, Sebastian Diecke.

#### Materials Availability

All cell lines generated in this study are potentially available on request to the lead contact. The availability depends on CITES and Nagoya protocol restrictions.

### EXPERIMENTAL MODEL AND SUBJECT DETAILS

#### Sumatran rhinoceros Kertam

The male Sumatran rhinoceros (SR, also known as hairy rhinoceros or Asian two-horned rhinoceros, Bornean subspecies, *Dicerorhinus sumatrensis harrisoni)*, Kertam (also called Tam) was captured from the wild in 2008, and subsequently housed in the Tabin Wildlife Reserve Sabah in Lahad Datu, Malaysia where he died in May 2019. All medical procedures were legalized by the Sabah Wildlife Department and performed by authorized veterinarians.

#### Human cell lines

Human iPS cells HMGUi-001-A (https://hpscreg.eu) were derived from fibroblasts of a healthy woman and were obtained from Heiko Lickert (Helmholtz Zentrum München, Munich, Germany). HMGUi-001-A hiPSCs were maintained in custom-made Essential 8 Medium [DMEM/F12 HEPES (Thermo Fisher, #11330032) supplemented with L-ascorbic acid 2-phosphate (Sigma, #A8960), insulin (CS Bio, #C9212-1G or Sigma, #91077C-1G), human transferrin (Sigma,#T3705-1G), sodium selenite (Sigma, #S5261-10G), bFGF (PeproTech, #100-18B), TGFβ1 (PeproTech, #100-21C) and sodium bicarbonate 7.5% solution (Fisher Scientific, #25080-094), according to (Chen et al., 2011)] on cell culture dishes coated with Geltrex (Thermo Fisher, #A1569601) at 37°C, 5% CO_2_, 5% O_2_ and passaged every 3-4 days using PBS 0.5 mM EDTA solution (PBS/EDTA, Thermo Fisher, #14190250 and #15575020) in ratios of 1:6 and 1:12. Cell line identity was authenticated by genotyping (STR analysis, GenePrint® 10 System, Promega Corporation) after banking.

### METHOD DETAILS

#### Derivation of SR primary fibroblast

The skin biopsy from the SR Kertam was taken under general anaesthesia in the *regio axillaris* during a general health and reproductive assessment. The biopsy area was widely disinfected with Octenisept Spray (TM, SCHÜLKE & MAYR GmbH, #121411) and the deep skin biopsy was achieved by using a biopsy punch device (KAI Medical, 4 mm in diameter, #BP-40F), sterile surgical forceps and a sterile scalpel. The recovered tissue was immediately transferred into cell culture medium [DMEM, high glucuose, GlutaMAX (Thermo Fisher, #10566016) supplemented with 10% fetal bovine serum (FBS, biowest, #S1400), 1X Penicillin-Streptomycin (biowest, #L0022), 1X Antibiotic-Antimycotic (biowest, #L0010)] and shipped at 4°C. The wound was sewed with simple surgical suture using 3/0 seam material with a sharp needle (Supramid®, 3/0 HS23 - 0,45m B. Braun Petzold, #C0712256). The suture site was covered against flies and better wound healing with veterinary aluminum wound spray (Pharmamedico GmbH, #03691157).

Upon arrival in the cell culture lab, the biopsy was diced into small pieces and cultured in DMEM/TCM 199 [1:1, DMEM, high glucuose, GlutaMAX, HEPES (Gibco, #32430027) : Medium 199, HEPES (Gibco, #22340020)] supplemented with 10% FBS and 2 ng/ml basic fibroblast growth factor (bFGF, Peprotech, #AF-100-18B) in humidified air at 38°C, 5% CO_2_ and 5% O_2_.

After expansion of cells in culture, they were subcultured every 4-6 days, frozen in DMEM/TCM 199 (1:1) with 20% FBS and 10% DMSO, and subsequently stored in liquid nitrogen.

#### Species analysis

To verify the identity of our SR primary fibroblasts, we applied a standardized species identification test published by Ewart et al., 2018. In brief, a cryovial containing primary fibroblasts was quickly thawed using a 37°C water bath. The cell suspension was transferred into a 1.5 ml Eppendorf tube and centrifuged at 300xg for 5 min at room temperature. The supernatant was aspired and the cell pellet washed once in 1 ml PBS. After centrifugation at 300xg for 5 min at room temperature, the cell pellet was resuspended in 300 μl PBS and genomic DNA (gDNA) automatedly extracted using the Maxwell RSC Blood DNA Kit (Promega, #AS1400) and a Maxwell RSC instrument according to the manufacturer’s instructions (quick protocol from point 2 on). gDNA was eluted in 85 μl and its concentration measured by nanodrop. Subsequently, PCR was performed in 50 μl reaction volume comprising 100 ng template DNA, 1 μl Phire Hot Start II DNA Polymerase (Thermo Fisher, #F-122S), 1X Phire Reaction buffer, 200 μM dNTP mix (Thermo Fisher #18427-013), 0.5 μM of each “universal rhino primers” (RID_FWD: 5′-AACATCCGTAAATCYCACCCA-3′and RID_REV: 5′-GGCAGATRAARAATATGGATGCT-3′) applying the following cycling protocol: 30 seconds initial denaturation at 98°C, 30 cycles of 5 seconds denaturation at 98°C, 5 seconds annealing at 55°C, 5 seconds extension at 72°C, and 1 minute final extension at 72°C. 9.6 μl of the PCR reaction was subjected to gel electrophoresis [2% TAE agarose gel, Roti-Gel Stain (Roth, #3865.2), 100 bp DNA ladder (NEB, #N3231L)] to validate exclusive amplification of the species specific 230 bp cytochrome-b region. The remaining PCR reaction was purified using the GeneJet PCR purification kit (Thermo Fisher, #K0702) according to the manufacturer’s instructions. The nucleotide sequence was confirmed by Sanger sequencing and alignment to the published sequence.

#### Proliferation analysis

To calculate the cell doubling rate, SR primary fibroblasts were plated on four 3×24-well-plates coated with attachment factor (3.0*10^4 cells/24-Well) in skin media. After 24, 48, 72, and 96 hours, one 3×24-well-plate was fixed each. To this end, medium was aspired, cells were washed once with PBS, and subsequently incubated for 15 minutes at room temperature in BD Cytofix solution (BD Biosciences, #554655). Afterwards, cells were washed three times and stored in PBS. Plates were sealed with parafilm and kept in the fridge in the dark until further use.

When all time points were collected, cells were incubated slowly shaking for 30 minutes at room temperature in staining solution [PBS with 0.1% Triton X-100 (Carl Roth, #3051.3), 2 drops/ml DAPI (NucBlue FIxed cell Stain Ready Probe, Thermo Fisher, #R37606)]. Subsequently, cells were washed three times with PBS. Per timepoint, 3 images (1 image per 24-well) were acquired with a LEICA DIMi8 microscope and the LAS X Software (10x magnification, 500 milliseconds exposure time). Single nuclei were automatedly counted using ImageJ software.

#### Generation of SR-iPSCs

After thawing, SR primary fibroblasts were maintained at 37°C and 5% CO_2_ on cell culture plates coated with attachment factor (Thermo Fisher, #S006100) in skin media [DMEM, high glucuose, GlutaMAX, pyruvate supplemented with 10% FBS, 1X MEM non-essential amino acids (NEAA, Thermo Fisher, #11140035) and 10 ng/ml bFGF]. With minor changes, reprogramming was performed as described previously for the northern white rhino (Hayashi et al. *submitted;* Korody et al., 2021). In brief, two days before transduction (day −2), fibroblasts were detached by incubation with TrypLE select enzyme (Thermo Fisher, #12563011) for 8 minutes at 37°C, and 2.4×10^4 cells were plated each in 3×12-wells coated with attachment factor in skin media. On the next day, medium was changed. On the day of transduction (day 0), cells of 1×12-well were detached with TrypLE and counted (~9.25×10^4 cells).

Based on the obtained cell number, the amount of Sendai viruses (CytoTune – iPS 2.0 Sendai Reprogramming Kit, Thermo Fisher, #A16517, Lot #L2160042) needed for transduction of cells in 1×12-well with an multiplicity of infection (MOI) of 10:10:6 (KOS:c-Myc:Klf4) was calculated. The medium of 2×12-wells was changed to skin medium containing 10 μg/ml polybrene (Merck, #TR-1003-G). Sendai virus mixture was added to 1×12-well. The other 1×12-well was not transduced and served as negative control. The plate was sealed with parafilm and centrifuged at room temperature and 800xg. After 20 minutes centrifugation, the plate was turned and centrifugated for another 10 minutes. Subsequently, parafilm was removed and the plate incubated overnight at 37°C and 5% CO_2_. On the next day (day 1), cells were washed once with skin medium and subsequently fed with skin medium. Skin medium was changed on day 2, day 4 and day 6. On day 7, transduced cells were carefully detached by incubation with TrypLE for 2-3 minutes at 37°C, and 5 different numbers of transduced cells (1.5, 3.0, 5.0, 7.5, 10.0 × 10^3) were plated on 5×6-wells [coated with Geltrex and laid with 2.0×10^5 mitomycin C-treated mouse embryonic fibroblasts (MEFs, tebu-bio, #MEF-MITC)] in skin medium. Additionally, 20×10^3 and 40×10^3 cells were plated feeder-free on Geltrex coated 2×6-wells. On the next day (day 8) medium of cells plated on MEFs was changed to KB medium [39% DMEM, high glucose (Thermo Fisher, #41965039), 39% complete FGM (Lonza, #CC3132), 20% knockout serum replacement (KSR, Thermo Fisher, #10828028), 1x NEAA, 1x GlutaMAX (Thermo Fisher, #35050061), 0.1 mM 2-Mercaptoethanol (Thermo Fisher, #21985023), and 12 ng/ml bFGF], whereas feeder-free cultures were changed to mTeSR1 medium (STEMCELL Technologies, #05850). From now on, medium was changed daily. On day 19, 8 and 4 single colonies were picked from KB on MEFs and feeder-free mTeSR1 wells, respectively. One promising clone observed in KB media on MEFs was split in 2 halves. One half was picked on day 19 (line #8), the other one on day 21 (line #9). Additionally, 2 and 5 individual colonies were isolated on this day from KB on MEFs and feeder-free mTeSR1 conditions, respectively. Depending on the previous culturing condition (KB or mTeSR1), picked single colonies were transferred to individual Geltrex coated 24-wells, which were either laid with 5.0×10^4 MEFs/well or kept feeder-free, respectively. The culturing media (KB or mTeSR1) was supplemented with 10 μM ROCK inhibitor (Y-27632, Selleck Chemicals, #SEL-S1049-10MM) at the day of picking and on the day after.

Three lines (#8, #9, #14; #8 and #9 were isolated from the same original clone) – all originating from KB on MEFs conditions - grew successfully and displayed nice morphology. The other lines could not be propagated further and were discarded. For splitting, cultures were incubated for 7-8 minutes at 37°C in PBS/EDTA. KB medium was supplemented with 10 μM ROCK inhibitor for 24 hours after splitting. Line #8, #9, #14 were cryopreserved in KSR, 10% Dimethyl Sulfoxid (DMSO, Sigma, #D22660). Subsequent experiments were performed with line #9.

#### Clearance of Sendai virus encoded RNA

To generate vector-free SR-iPSC lines, we picked 12 subclones from line #9 at passage 4 on individual Geltrex coated 24-wells laid with 5.0×10^4 MEFs/well. KB medium was supplemented with 10 μM ROCK inhibitor for 24 hours after picking. All subclones were expanded and samples for RNA extraction collected at passages 7, 8 and 9. RNA was isolated using the RNeasy Mini Kit (Qiagen, #74106) according to the manufacturer’s instructions. RNA was eluted in 40 μl water and its concentration measured by Nanodrop. 3 μl of each sample were reversely transcribed with SuperScript III First-Strand Synthesis SuperMix (Thermo Fisher, #18080400) using oligo(dT) primer and following the corresponding protocol of the manufacturer. cDNA was diluted 1:5 with water and 2 μl used for PCR [20 μl reaction volume, DreamTaq Green PCR Master Mix (Thermo Scientific, #K1081), 0.5 μM each forward and reverse primer, primer sequences provided in the manual of the CytoTune – iPS 2.0 Sendai Reprogramming Kit (Thermo Fisher, #A16517); PCR program: 5 minutes initial denaturation at 94°C, 35 cycles of 30 seconds denaturation at 95°C, 30 seconds annealing at 55°C, 30 seconds elongation at 72°C, followed by final extension for 7 minutes at 72°C]. Absence of sendai-virus encoded genes (SeV, KOS, Klf4, c-Myc) was confirmed by gel electrophoresis (2% TAE agarose gel, Roti-Gel Stain, 100 bp DNA ladder) for subclones #9A, #9C and #9K. Subsequent experiments were performed with line #9K.

#### SR-iPSC maintenance culture

SR-iPSCs were cultured on Geltrex or attachment factor coated cell culture dishes laid with MEFs (21-29 cells/cm^2^) at 37°C and 5% CO_2_. When cells reached 80-90% confluency, they were split with PBS/EDTA (6-7 minutes at 37°C) in varying ratios (1:3-1:12). ROCK inhibitor was added to the medium at 5 μM for the first 24 hours after passaging.

#### Karyotyping of SR-iPSCs

To check the genomic integrity of SR-fibroblasts and SR-iPSCs, Giemsa-trypsin-Giemsa (GTG, G-banding) banding technique was performed as described in Hayashi et al. *submitted* (according to Seabright, 1971; Weise and Liehr, 2009). Additionally, centromeric heterochromatin staining (CBG, C-banding) was performed following a protocol adapted from (Salamanca and Armendares, 1974). The methods are briefly summarized below.

##### Chromosome preparation

SR-fibroblasts were split with TrypLE (incubation for 8 minutes at 37°C) and 2.5*10^5 cells were plated each in two T25flasks coated with attachment factor. Cells were fed with skin media, which was changed every other day. SR-iPSCs were detached with PBS/EDTA for 6-7 minutes at 37°C and 6.25*10^5 cells were plated each in two Geltrex coated T25flasks in cKB supplemented with 5 μM ROCK inhibitor, which was changed daily.

At 50% confluency, cells were treated with 0.1 g/ml colcemide (Biochrom AG, #L6221) for 2.5 hours at 37°C, and subsequently harvested using Trypsin-EDTA (Biochrom AG, # L2143). Enzymatic digestion was stopped by adding medium containing 10% KSR. Cells from two T25 flasks were pooled in one 50 ml Falcon tube and centrifuged at room temperature for 5-10 minutes at 450xg. The supernatant was discarded except for 1 ml, which was used to resuspend the cell pellet and to transfer it into a 15 ml falcon tube. Cells were treated with 10 ml hypotonic 0.075 M KCl solution (Merck, #1049360500) for 20 minutes at 37°C, followed by adding ~1 ml freshly prepared ice-cold fixative (Methanol / Acetic acid, ratio 3:1, Merck, #1060092500 and #1000632500) and careful mixing. After centrifugation at room temperature for 5-10 minutes at 450xg, the supernatant was discarded except for ~1 ml to resuspend the cell pellet. Subsequently, cells were washed three times with 5-10 ml fixative, centrifuged and finally resuspended in 2 ml fixative and stored at −20°C. Before slides were prepared, the cell pellet was washed again and resuspended in fresh fixative. The cell suspension was dropped on a cleaned, cooled and wet glass slide, air dried and baked over night at 70°C for subsequent GTG and CBG banding.

##### GTG banding

Slides were processed in a glass cuvette at room temperature with 1 ml trypsin stock solution (Gibco, #27250-018, 6 g in 100 ml in PBS Dulbecco, Biochrom, # L1825) diluted in 100 ml PBS for 2 seconds, put in a glass cuvette with 150 ml buffer pH 6,88 (Merck, # 1072941000) and transferred to a second glass cuvette with 150 ml buffer pH 6,88. Subsequently, slides were stained in a glass cuvette with Giemsa solution [6 ml Giemsa stain (Sigma, # GS500) and 75 ml puffer pH 6,88) for 3 minutes, rinsed with aqua deion. and finally air dried. Metaphase chromosomes were automatically scanned and imaged with the Zeiss Axio.Z2 microscope using the scanning system Metafer and the Ikaros V5.1 software (both Metasystems). The average resolution was ~200 bands per haploid chromosome set. In total, 10-20 metaphase spreads were analyzed per cell line.

##### CBG banding

For CBG, slides were treated with 0.2 N hydrochloric acid (Merck, # 109057) for 1 hour at room temperature, rinsed with aqua deion., incubated for 7 minutes in filtered 5% barium hydroxide solution (Merck, #101737), rinsed again with aqua deion., left for 5 minutes in 0.2 N hydrochloric acid followed by 1 hour incubation in 2xSSC (Invitrogen, # 15557-036) at 60°C, rinsed with aqua deion., stained with Giemsa stain solution for 20 minutes, finally rinsed with aqua deion. and air dried. Ten metaphases were captured under an Axioskop 20 light microscope (Zeiss) at 100x oil immersion objective and analyzed by IKAROS V5.1 software (Metasystems).

#### Three germ layer differentiation potential

Endoderm was differentiated from SR-iPSCs using the StemMACS Trilineage EndoDiff medium (Miltenyi Biotec, #30-115-659) following the manufacturer’s instructions. In brief, SR-iPSCs were detached with PBS/EDTA and resuspended in 1 ml MEF conditioned KB medium (cKB) supplemented with 10 μM ROCK inhibitor. cKB was generated by incubating MEFs overnight in KB medium (2 ml per 2×10^5MEFs/6-well). 2-Mercaptoethanol and FGF were added freshly before using the medium on SR-iPSCs. The cell suspension was filtered through a 70 μm strainer (Miltenyi Biotec, #130-095-823) and single living cells counted using Trypan Blue (Thermo Fisher, #T10282) and the CellDrop Bright Field Cell Counter (Biozym, # 31CELLDROPBF-UNLTD). Subsequently, 1.25×10^5 cells were plated per Geltrex coated 24-well (containing cover slip) in 0.5 ml cKB medium supplemented with 10 μM ROCK inhibitor (day 0). On day 1, 0.5 ml cKB medium without additional ROCK inhibitor was added. On day 2, medium was changed to EndoDiff medium. On day 3-6, EndoDiff medium was changed daily. On day 7, cells were washed twice in PBS+/+ (PBS, calcium, magnesium; Thermo Fisher, #14040141), and subsequently fixed for 15 minutes at room temperature in BD cytofix solution (BD, #554655). After washing twice in PBS+/+, fixed cells were stored at 4°C until further use.

To represent the mesoderm, SR-iPSCs were differentiated into cardiomyoctes using a PSC cardiomyocyte differentiation kit (Thermo Fisher, #A2921201) according to the manufacturer’s instructions. In brief, SR-iPSCs were detached with PBS/EDTA and plated in cKB supplemented with 1X RevitaCell (Thermo Fisher, #A2644501) in various ratios (corresponding to 1:4-1:12) in Geltrex coated 12-wells. For the next two days, cells were fed daily with cKB without RevitaCell. Day 3 after plating, medium was changed to Cardiomyocyte Differentiation Medium A (day 0). On day 2, the medium was slowly aspired and carefully 2 ml pre-warmed Cardiomyocyte Differentiation Medium B was added per well. On day 4, cultures were cautiously shaken to swirl up dead cells, medium was removed, and 2 ml pre-warmed Cardiomyocyte Maintenance Medium added per well. From now on, Cardiomyocyte Maintenance Medium was changed every other day. Clusters of beating cells were observed from day 9 on. Around day 11, cultures were changed to RPMI B-27 medium [RPMI 1640 (Thermo Fisher, #11875093) supplemented with B-27 (Thermo Fisher, #17504044)], which was changed every other day. After day 15, cardiomyocytes were fixed as described before and stored at 4°C until further use.

The neural (ectodermal) differentiation of SR-iPSCs was induced by a modified version of the dual SMAD inhibition protocol published by Chambers et al., 2009. In brief, SR-iPSC were detached with PBS/EDTA and 6.0×10^5 cells plated per 6-well in cKB supplemented with 5 μM ROCK inhibitor. On the next day, medium was changed to cKB without ROCK inhibitor. The day after (day 0), medium was changed to neural induction medium [NIM; DMEM/F12 (Thermo Fisher, #12660012), 1x B-27, 1x N-2 (Thermo Fisher, #17502048), 10 μM SB431542 (Reagents Direct, #21-A94), 2 μM Dorsomorphin (Biovision, #1686-5)]. NIM was exchanged daily for 5 consecutive days. On day 6, cells were washed once with PBS and subsequently detached using Accutase (Thermo Fisher, #A1110501). Enzymatic reaction was stopped after 8 minutes by adding neural expansion medium [NEM; 0.5X advanced DMEM/F12 (Thermo Fisher, #12634010), 0.5X Neurobasal Medium (Thermo Fisher, #21103049); 2X N-2 (Thermo Fisher, #17502048)] supplemented with 5 μM ROCK inhibitor. Cells were detached using a cell scraper and passed through a 70 μm strainer and subsequently centrifuged at 300xg for 3 minutes at room temperature. The supernatant was discarded, cells resuspended in NEM supplemented with 5 μM ROCK inhibitor, counted, and 1×10^5 cells plated per Geltrex coated 24-well (containing cover slip). Medium was changed every day with NEM. On day 12, cells were fixed as described previously and stored at 4°C until further use.

#### Cerebral organoids

Cerebral organoids with mainly cortical identity were generated from SR-iPSCs with the STEMdiff™ Cerebral Organoid Kit (STEMCELL Technologies, #08570) based on a previously described protocol (Lancaster and Knoblich, 2014) with some modifications during EB formation (days 0-5). Briefly, SR-iPSC grown in clusters on MEFs were detached using ReleSR (STEMCELL Technologies, #05872) according to the manufacturer’s instructions. Detached cell aggregates of 50 – 200 μm size were transferred to a 15 ml tube using a 5 ml serological pipette. After settling down, medium was removed, aggregates resuspended in 2ml KB medium, plated in one well of a 6-well plate with ultra-low attachment surface (Corning, #3471) and kept at 37°C and 5% CO_2_. KB medium was changed on days 1 and 3 by transferring the EBs to a 15ml tube as described above. From day 5 on, the timing and the media of the organoid kit were used, with exception of day 7, where 2% Matrigel (Corning, # 356231) was added to the medium for neuroepithelial expansion, instead of droplet embedding of single EBs. On day 10, the medium was exchanged for maturation medium and organoids were placed on an orbital shaker rotating at 65rpm in the incubator from there on.

#### Cryosectioning of cerebral organoids

Organoids between 1 and 3 months of age were fixed in 4% PFA (Affimetrix, #199431LT) for 30 minutes at room temperature, washed 3 times in PBS and left overnight at 4°C in 30% sucrose solution. Next day, organoids were embedded in warmed 7.5% gelatin solution (VWR, # 24350.262), cooled down on ice, snap frozen in isopentane/dry ice (−50°C) and stored at −80°C, according to Lancaster et al. 2014. Organoids were cryosectioned in 20 μm thick slices using a MicroM HM 560 Cryostat (Thermo Fisher), collected on Superfrost Ultraplus glass slides (Roth, # H867.1) and stored at −80°C. For immunohistochemical analysis, the gelatin was removed from the slides by incubation in PBS at 42°C for 20 minutes. Slides were blocked in NDS buffer and immunostaining performed as described in the next section.

#### Immunofluorescence

To stain SR-iPSCs in pluripotent state or after 2D-differentiation into cells of the three germ layers, cells were fixed as described previously. PBS+/+ was aspired and cells blocked for 1 hour shaking at room temperature either in 1xPBS+/+, 5% normal goat serum (NGS; Abcam, #ab7481) for surface markers, or for intracellular and nuclear markers in blocking buffer [PBS+/+, 0.2% BSA (Biomol, # 1400100), 0.3 % Triton X (Sigma, # T8787), 10 x NGS or normal donkey serum (NDS, abcam, #ab7475)]. Afterwards, primary antibodies were diluted as indicated in Table 1 in 1xPBS+/+, 1% BSA for surface markers or blocking buffer for intracellular and nuclear markers. After incubation shaking at 4°C overnight, cells were washed three times à 15 minutes in PBS+/+. Secondary antibodies were diluted in PBS+/+ containing DAPI, and cells incubated in the solution for ~2 hours shaking at room temperature. Subsequently, cells were washed three times in PBS+/+. Cells grown on cover slips were mounted on slides using Fluoromount-G (Thermo Fisher, #00-4958-02). Images were captured and processed with a Leica DMi8 microscope and the LASX software (including THUNDER computational clearing method).

**Table 1.**
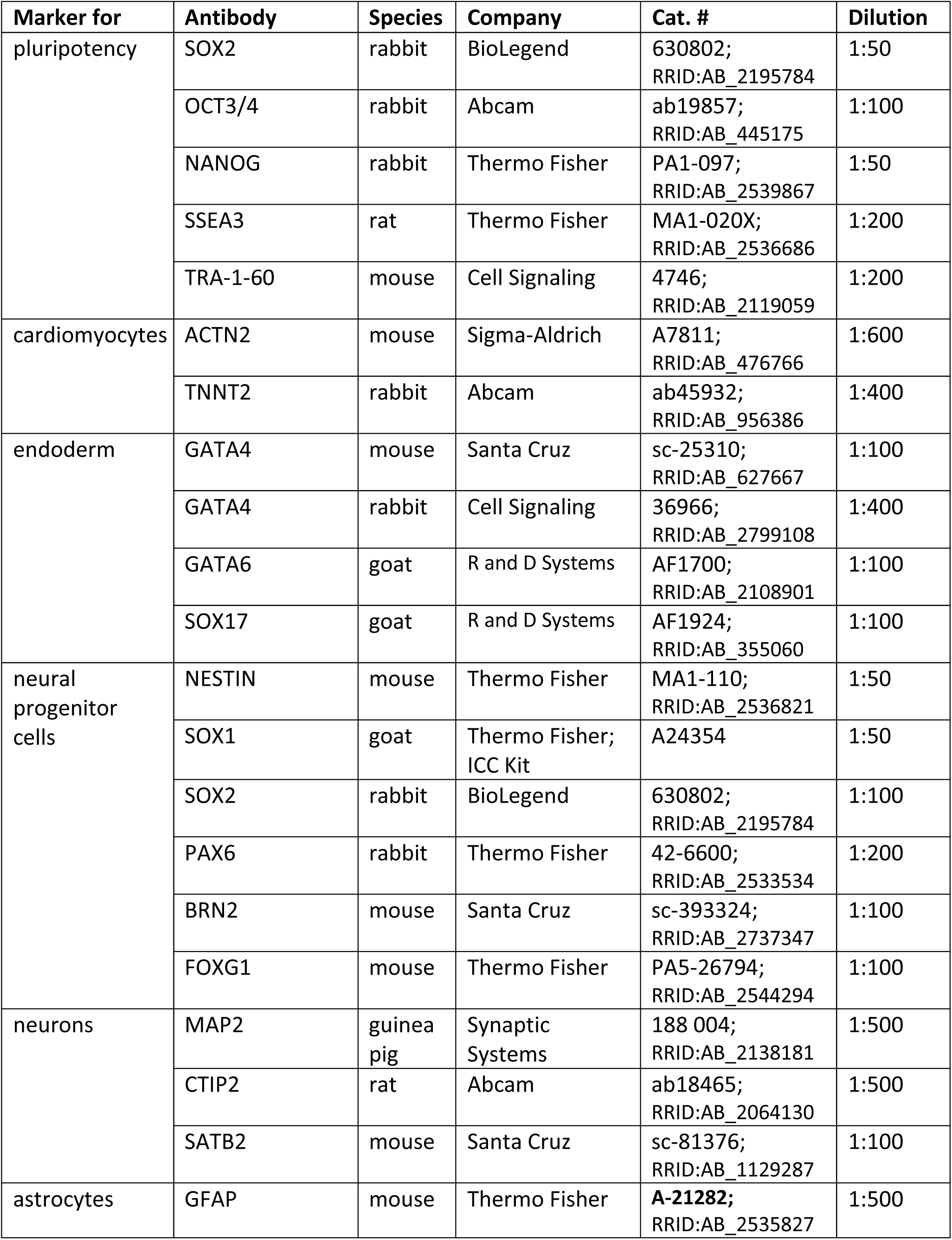
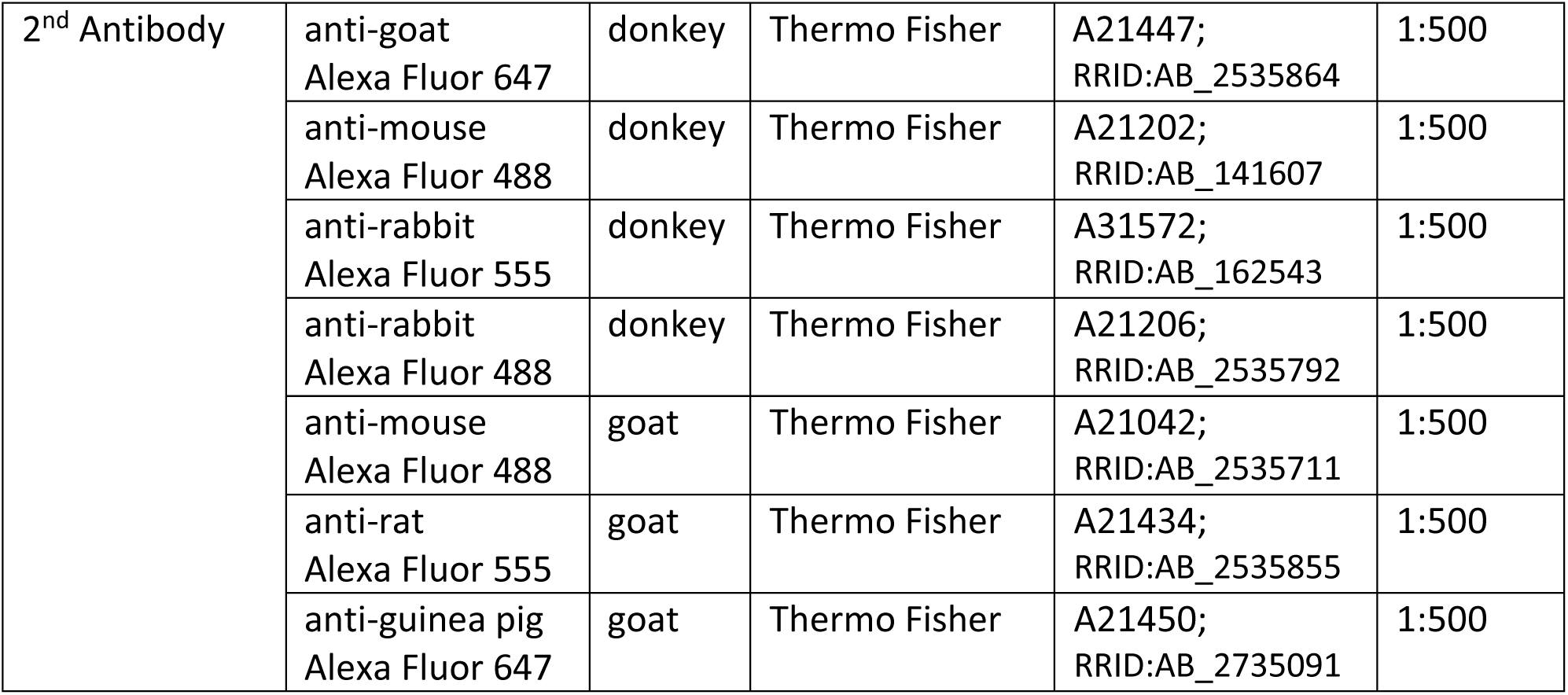
List of antibodies.

| **Marker for** | **Antibody** | **Species** | **Company** | **Cat. #** | **Dilution** |
| --- | --- | --- | --- | --- | --- |
| pluripotency | SOX2 | rabbit | BioLegend | 630802;<br>RRID:AB_2195784 | 1:50 |
|  | OCT3/4 | rabbit | Abcam | ab19857;<br>RRID:AB_445175 | 1:100 |
|  | NANOG | rabbit | Thermo Fisher | PA1-097;<br>RRID:AB_2539867 | 1:50 |
|  | SSEA3 | rat | Thermo Fisher | MA1-020X;<br>RRID:AB_2536686 | 1:200 |
|  | TRA-1-60 | mouse | Cell Signaling | 4746;<br>RRID:AB_2119059 | 1:200 |
| cardiomyocytes | ACTN2 | mouse | Sigma-Aldrich | A7811;<br>RRID:AB_476766 | 1:600 |
|  | TNNT2 | rabbit | Abcam | ab45932;<br>RRID:AB_956386 | 1:400 |
| endoderm | GATA4 | mouse | Santa Cruz | sc-25310;<br>RRID:AB_627667 | 1:100 |
|  | GATA4 | rabbit | Cell Signaling | 36966;<br>RRID:AB_2799108 | 1:400 |
|  | GATA6 | goat | R and D Systems | AF1700;<br>RRID:AB_2108901 | 1:100 |
|  | SOX17 | goat | R and D Systems | AF1924;<br>RRID:AB_355060 | 1:100 |
| neural progenitor cells | NESTIN | mouse | Thermo Fisher | MA1-110;<br>RRID:AB_2536821 | 1:50 |
|  | SOX1 | goat | Thermo Fisher;<br>ICC Kit | A24354 | 1:50 |
|  | SOX2 | rabbit | BioLegend | 630802;<br>RRID:AB_2195784 | 1:100 |
|  | PAX6 | rabbit | Thermo Fisher | 42-6600;<br>RRID:AB_2533534 | 1:200 |
|  | BRN2 | mouse | Santa Cruz | sc-393324;<br>RRID:AB_2737347 | 1:100 |
|  | FOXG1 | mouse | Thermo Fisher | PA5-26794;<br>RRID:AB_2544294 | 1:100 |
| neurons | MAP2 | guinea pig | Synaptic Systems | 188 004;<br>RRID:AB_2138181 | 1:500 |
|  | CTIP2 | rat | Abcam | ab18465;<br>RRID:AB_2064130 | 1:500 |
|  | SATB2 | mouse | Santa Cruz | sc-81376;<br>RRID:AB_1129287 | 1:100 |
| astrocytes | GFAP | mouse | Thermo Fisher | **A-21282**;<br>RRID:AB_2535827 | 1:500 |
| 2<sup>nd</sup> Antibody | anti-goat<br>Alexa Fluor 647 | donkey | Thermo Fisher | A21447;<br>RRID:AB_2535864 | 1:500 |
|  | anti-mouse<br>Alexa Fluor 488 | donkey | Thermo Fisher | A21202;<br>RRID:AB_141607 | 1:500 |
|  | anti-rabbit<br>Alexa Fluor 555 | donkey | Thermo Fisher | A31572;<br>RRID:AB_162543 | 1:500 |
|  | anti-rabbit<br>Alexa Fluor 488 | donkey | Thermo Fisher | A21206;<br>RRID:AB_2535792 | 1:500 |
|  | anti-mouse<br>Alexa Fluor 488 | goat | Thermo Fisher | A21042;<br>RRID:AB_2535711 | 1:500 |
|  | anti-rat<br>Alexa Fluor 555 | goat | Thermo Fisher | A21434;<br>RRID:AB_2535855 | 1:500 |
|  | anti-guinea pig<br>Alexa Fluor 647 | goat | Thermo Fisher | A21450;<br>RRID:AB_2735091 | 1:500 |

## ACKNOWLEDGMENTS

We thank our Malayan colleagues Dr John Payne und Dr Zainal Zahari Zainuddin for the fruitful collaboration, and our colleagues from Diecke lab for technical assistance. This work was supported in part by grants from the Malayan NGO Sime Darby Foundation and the German government BMBF 01LC1902B BioRescue.

## AUTHOR CONTRIBUTIONS

Conceptualization, T.B.H., S.D., C.G., and M.D.; Validation, V.Z., S.F., and A.W., Methodology, V.Z. and S.F.; Investigation, V.Z., S.F., N.K., A.W. and S.C.; Resources, F.G., R.H., S.H. and T.B.H.; Writing – Original Draft, V.Z. and S.F., Writing –Review & Editing, V.Z., S.F., T.B.H., S.H., and S.D., Funding Acquisition, T.H.B. and S.D.

## SUPPLEMENTAL INFORMATION

**Supplementary Figure 1.**
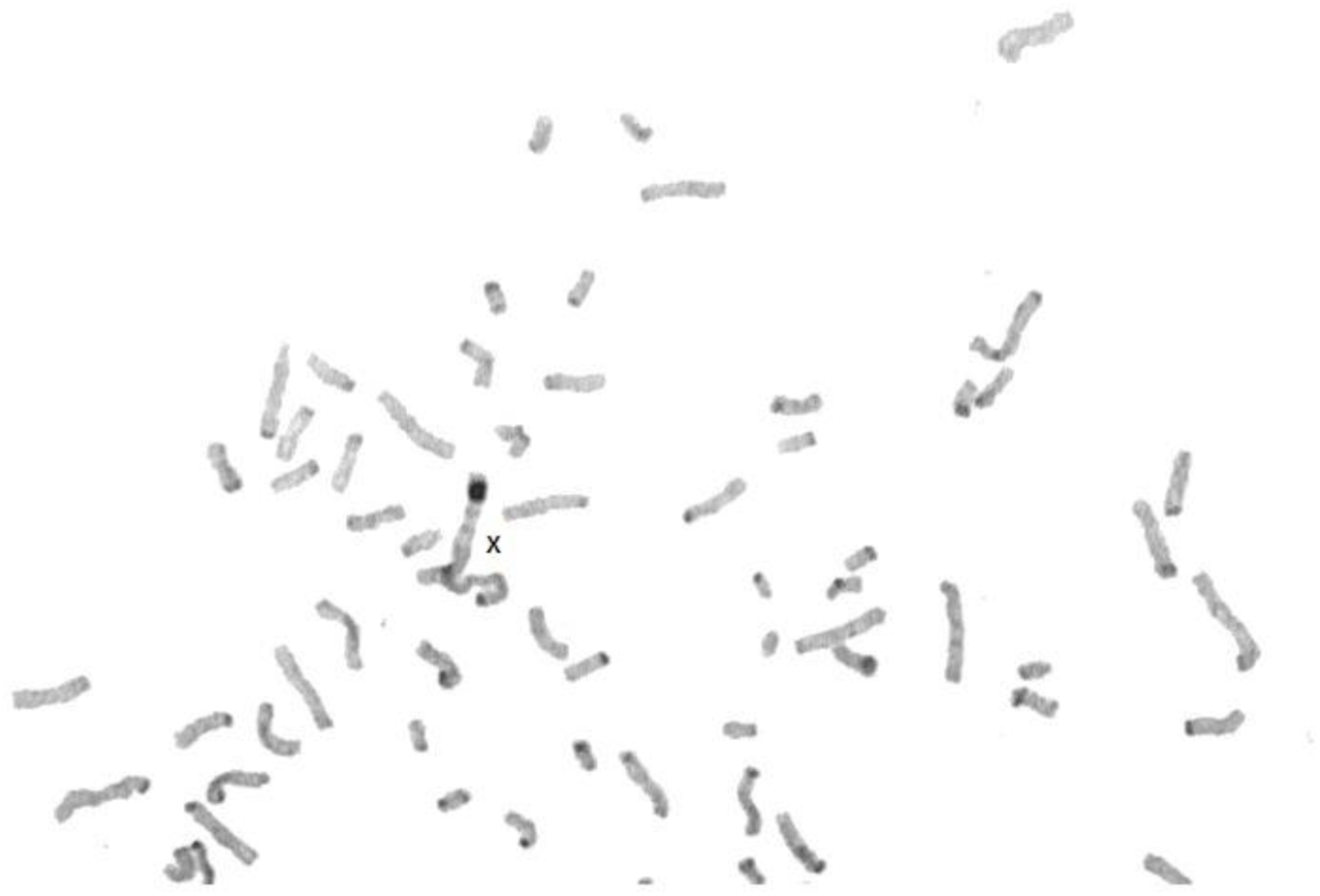
CTG banding of SR-iPSCs. confirms large block of heterochromatin at the terminal end of the long (q) arm of the X chromosome. Related to Figures 1E and 1F.

**Supplementary Figure 2.**
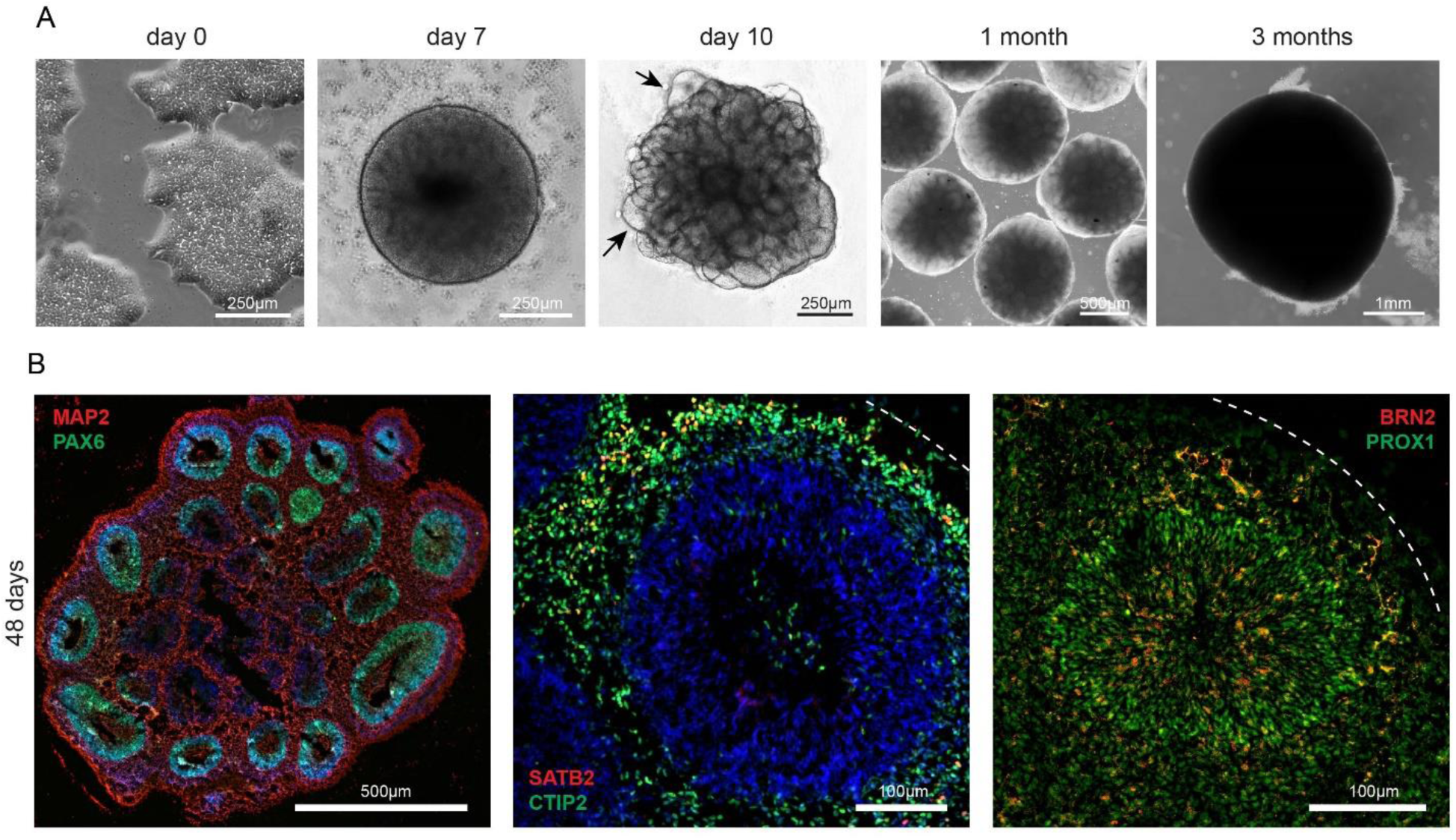
Generation of human cerebral organoids. **A**. Exemplary phase contrast images during organoid generation from human iPSCs at corresponding days of differentiation. Day 0: Feeder-free human iPSC colonies before dissociation; day 7: embryoid body showing a smooth and brightened surface; day 10: after addition of matrigel to the medium, organoids showed neuroepithelial bud outgrowth resembling neural tube like structures (arrows), which are still visible after 1 month in culture. 3 months: mature organoid. Scale bars are defined within each image. **B**. Immunostaining of 48 days old human brain organoids. Radially organized neural progenitors stain positive for PAX6, while neurons express MAP2, CTIP2, BRN2 and PROX1. Scale bars are defined within each image.

